# Affectively biased competition: sustained attention is tuned to rewarding expressions and is not modulated by norepinephrine receptor gene variant

**DOI:** 10.1101/442863

**Authors:** Kevin H. Roberts, Maria G. M. Manaligod, Colin J. D. Ross, Daniel J. Müller, Matthias J. Wieser, Rebecca M. Todd

**Author notes:** Address correspondence to: Rebecca Todd, PhD, Department of Psychology, University of British Columbia, 2136 West Mall, Vancouver, BC, V6T 1Z4, Canada.

## Abstract

It is well established that emotionally salient stimuli evoke greater visual cortex activation than neutral ones, and can distract attention from competing tasks. Yet less is known about underlying neurobiological processes. As a proxy of population level *biased competition*, EEG steady-state visual evoked potentials are sensitive to competition effects from salient stimuli. Here we wished to examine whether individual differences in norepinephrine activity play a role in emotionally-biased competition.

Our previous research has found robust effects of a common variation in the *ADRA2B* gene, coding for alpha2B norepinephrine (NE) receptors, on emotional modulation of attention and memory. In the present study, EEG was collected while 87 carriers of the *ADRA2B* deletion variant and 95 non-carriers (final sample) performed a change detection task in which target gratings (gabor patches) were superimposed directly over angry, happy, and neutral faces. Participants indicated the number of phase changes (0-3) in the target. Overlapping targets and distractors were flickered at a distinct driving frequencies. Relative EEG power for faces vs. targets at the driving frequency served as an index of cortical resources allocated to each of the competing stimuli. Deletion carriers and non-carriers were randomly assigned to Discovery and Replication samples and reliability of results across samples was assessed before the groups were combined for greater power.

Overall happy faces evoked higher competition than angry or neutral faces; however, we observed no hypothesized effects of *ADRA2B*. Increased competition from happy faces was not due to the effect of low-level visual features or individuals low in social anxiety. Our results indicate that emotionally biased competition during sustained attention, while reliably observed in young adults, is not influenced by commonly observed individual differences linked to NE receptor function. They further indicate an overall pattern of affectively-biased competition for happy faces, which we interpret in relation to previously observed boundary conditions.

## Introduction

It is well established that emotionally relevant events are prioritized over less relevant ones, evoke higher levels of visual cortex activation, and are perceived as being relatively more vivid (Markovic, Anderson, & Todd, 2014; Pourtois, Schettino, & Vuilleumier, 2013; Todd, Talmi, Schmitz, Susskind, & Anderson, 2012). Emotionally relevant stimuli can also capture neurocognitive resources when in competition with explicit goals demanded by an experimental task [for review see (Carretie, 2014)]. Yet although biased attention to what is emotionally or motivationally relevant is ubiquitous, habitual tendencies to preferentially attend positively or negatively valenced information vary between individuals. Affective biases toward prioritizing negative or positive stimuli have been associated with personality traits such as neuroticism and extraversion respectively (Derryberry & Reed, 1994). When rigid or extreme, valenced biases are also associated with etiology and maintenance of psychopathology. For example, anxiety is associated with a bias to attend threat, and depression and addiction with reduced or enhanced attention to rewarding information, respectively (B. A. Anderson, 2016; Bar-Haim, Lamy, Pergamin, Bakermans-Kranenburg, & van, 2007; Dalgleish et al., 2003; Peckham, McHugh, & Otto, 2010; Surguladze et al., 2004).

Research using functional magnetic resonance imaging (fMRI) has provided evidence that such biases are associated with large-scale patterns of brain activation involving amygdala and striatal systems [for review see (Todd & Manaligod, 2017)]. Less is known about underlying neuronal and neuromodulator processes. Recent research has begun to employ EEG steady-state visual evoked potentials (ssVEPs) to investigate population-level neuronal processes underlying rapid visual cortex plasticity with aversive conditioning (Kastner-Dorn, Andreatta, Pauli, & Wieser, 2018; McTeague, Gruss, & Keil, 2015; Miskovic & Keil, 2013; Wieser, Reicherts, Juravle, & von Leupoldt, 2016). As a proxy of population level *biased competition*, ssVEPs measured at the scalp can be sensitive to competition effects from salient stimuli (Wieser, McTeague, & Keil, 2012). When two spatially overlapping stimuli are driven at distinct temporal frequencies, power at the driving frequency of each can be compared to index relative allocation of attention (Y. J. Kim, Grabowecky, Paller, Muthu, & Suzuki, 2007). A difference score can then be calculated as a measure of the degree to which a distractor captures resources from a task-relevant stimulus. Previous research has reported that higher levels of social anxiety are associated with biased competition for angry – relative to happy faces – when task-irrelevant emotional faces are overlaid with task-relevant gratings. For those with low social anxiety the reverse was true (Wieser et al., 2012).

The influence of emotional salience on selective attention is also modulated by the noradrenergic system, which influences gating and tuning of visual cortex neurons. Neurons in the locus coeruleus, the brainstem nucleus that primarily produces norepinephrine (NE), responds to both direct reward and punishment and very rapid learning of associations with both (Markovic et al., 2014; Sara, 2009; Sara & Bouret, 2012). A prominent model of arousal-biased cognition proposes that LC/NE activity serves to amplify neuronal responses in “local hotspots” to facilitate biased competitive processing of events that are relevant to current goals (Mather, Clewett, Sakaki, & Harley, 2016). NE modulation of excitation/inhibition in response to incoming sensory information allows for neural circuits to respond with optimal efficiency to the most salient information – regardless of valence (Berridge & Waterhouse, 2003). With regard to vision, rodent studies have found alpha2 receptors to mediate modulation of signal to noise ratio in primary visual cortex and can alter selectivity and tuning of visual cortex cells. NE activity can influence receptive field properties, such as alteration of feature selectivity (e.g., direction), influencing visual tuning in a context-dependent manner (Waterhouse & Navarra, 2018). A recent study pharmacologically manipulated NE levels in humans to find higher NE levels increased detection sensitivity and discrimination accuracy and increased consistency from trial to trial in evoked visual cortex EEG activity, showing a direct causal role for NE in boosting human visual cortex activity (Gelbard-Sagiv, Magidov, Sharon, Hendler, & Nir, 2018); however, whether NE influences electrophysiological markers of underlying neuronal mechanisms such as biased competition is still an open question.

As an index of individual differences in NE activity, a common deletion variant in the *ADRA2B* gene, which codes for inhibitory alpha2b NE receptors, is thought to be associated with higher levels of NE availability (de Quervain et al., 2007). The DEL301-303 polymorphism is a 9 bp in-frame deletion beginning at nucleotide 091, which results in the loss of three glutamic acid residues in the third intracellular loop of the receptor, which are important for phosphorylation [see (Small, Brown, Forbes, & Liggett, 2001)]. While studies directly examining the relationship between *ADRA2B* and NE availability are still lacking in humans, there is convergent evidence for supporting such a relationship. An examination of effects of the deletion variant in vitro indicated that the deletion variant decreased phosphorylation. This completely eliminated the short-term agonist-promoted desensitization of the receptor (Small et al., 2001). That is, it reduced the short term inhibitory function of the autoreceptor. A study that genetically engineered mice to impair activity of all three alpha2 receptor subtypes independently found that only alpha2b deficient mice had significantly higher levels of NE than wild-type controls (Makaritsis, Johns, Gavras, & Gavras, 2000). Thus, both of these studies show direct effects of genetic variation in alpha2B autoreceptor activity to be linked to increased NE availability. It has further been observed that effects of the deletion on behavior are equivalent to effects of administering yohimbine, an alpha2 noradrenergic antagonist, (de Quervain & Papassotiropoulos, 2006), indirectly supporting the hypothesis that this relationship is also observed in humans.

Our own research has found that carriers of the deletion variant show enhancement of emotional modulation of cognition relative to non-carriers: Deletion carriers show attentional prioritization of emotionally arousing words, enhanced neural and behavioral indices of perceptual vividness for emotionally relevant images, and stronger links between subjectively rated arousal during picture viewing and subsequent emotional enhancement of memory for the pictures (Todd et al., 2015; Todd et al., 2013; Todd et al., 2014). While findings have not been entirely consistent, a recent meta-analysis (Xie, Cappiello, Meng, Rosenthal, & Zhang, 2018) found reliable effects of carrying the deletion variant on emotional enhancement of attention, perception and memory. These effects were modulated by the specific cognitive processes indexed by the task, and were overall strongest in tasks indexing attentional and perceptual processes (Xie et al., 2018). With regard to effects of the deletion variant on sensitivity to emotional valence, some studies have found deletion carriers to prioritize emotionally arousing images in general (Todd et al., 2015), whereas others have found deletion carriers to show greater biases than non-carriers towards negative (Todd et al., 2013) or positive (Ehlers & Todd, 2018; Fairfield et al., 2019) stimuli. Our own studies have found that carriers of the deletion variant show a stronger bias to whatever stimuli are found to be more salient in the population in general in a given context — and whether positive or negative valence is more salient has varied with context. Outstanding questions concern boundary conditions of *ADRA2B* effects on emotionally modulated cognition, as specific cognitive processes are differentially modulated by emotion, and the distinct neurocognitive mechanisms underlying these effects.

In the biased attention by norepinephrine (BANE) model, we previously put forward a theoretical framework in which we hypothesized that increased NE availability putatively associated with carrying the *ADRA2B* deletion variant would bias attention by enhancing biased competition processes (Markovic et al., 2014). Thus, neuronal populations sensitive to features of affectively salient stimuli would increase firing and neuronal populations sensitive to competing features would be suppressed. The present study set out to directly test this hypothesis. As pharmacological interventions cannot precisely target specific NE receptors, genotyping for the deletion variant serves as a unique tool for examining individual differences in the NE systems mediated by alpha2B receptors. The relatively even distribution of the *ADRA2B* deletion variant the population allows for a natural experiment on individual differences linked to NE function on mechanistic neural processes underlying affectively biased attention.

To tap neuronal indices of affectively biased competition, we employed an experimental design designed by Wieser and colleagues (2012) using spatially overlapping neutral targets and emotional distractors to measure biased competition processes using ssVEPs (Wieser et al., 2012). Using this task, the authors previously found distinct patterns of biased competition to valenced facial expressions, with biased competition favoring positive expressions in participants with low social anxiety and to negative expressions in those with high social anxiety (Wieser et al., 2012). Thus, by overlaying task relevant and emotionally salient stimuli, this task provided a window into biased competition processes underlying individual differences in affectively biased attention. As recently reported, (Hedge, Powell, & Sumner, 2018), tasks that show reliable overall effects in general (e.g., consistent effects of emotional valence ACROSS individuals) are often insensitive to individual differences (e.g., individual differences in biases in sensitivity to valence). We reasoned that a task eliciting divergent patterns of biased competition linked to anxiety should also be sensitive to individual differences linked to putative NE availability. If *ADRA2B* genotype influenced biased competition at the level of neuronal populations, it would provide a level of mechanistic insight into affectively biased competition — indicating that NE modulates neuronal competition processes that give rise to affective biases — that is relatively rare and difficult to achieve in human studies. We hypothesized that *ADRA2B* deletion carriers would show higher levels of competition associated with attentional bias for emotionally expressive faces.

As previous studies had only investigated patterns of biased competition in populations either high or low in social anxiety (Wieser et al., 2012), a secondary question concerned whether we would observe overall valenced patterns of emotional modulation of attention in a relatively large sample of healthy young adults. Finally, we wished to examine whether we would replicate previously observed effects of social anxiety in a larger sample.

## Methods

### Participants

A total of 310 participants (76 males, 1 non-binary) were recruited from the University of British Columbia and received course credit or $20 for participating in the study. Participants were of European Caucasian descent, had no history of anxiety disorders, traumatic brain injury and epilepsy, and reported normal to corrected-to-normal vision. 113 were excluded due to a programming error in resulting in slow monitor refresh rates, such that no peaks were observable at the driving frequencies, 12 participants were excluded due to insufficient SNR among posterior electrodes, one participant was excluded after visual inspection of frequency amplitudes, and genotyping failed for two participants. There were additional errors in recording accuracy data for four participants, who were omitted from the analysis (due to a bug in the software, accuracy values were uninterpretable – not 0 or 1). Thus, the final N was 182 (46 males, 1 non-binary, mean age = 20.75 years, SD = 2.85, 87 ADRA2B deletion carriers). All procedures were approved by the clinical research ethics review board of the University of British Columbia. The sample size was determined a priori by power analysis based on effect sizes observed for *ADRA2B* effects on emotional bias in previous studies. After discovering our screen refresh error we collected additional data to meet our sample size estimates. To mitigate against spurious false positives found in targeted polymorphisms studies with relatively small sample sizes (Grabitz et al., 2018), we adopted a split-sample approach to the data analysis: Following data collection, equal numbers of *ADRA2B* deletion carriers and noncarriers were randomly assigned to the discovery and replication samples. There were 90 participants in the discovery sample (19 male, 1 non-binary, mean age = 20.71 *SD* = 2.74, 43 deletion carriers), and 92 participants in the replication sample (26 male, mean age = 20.79, *SD* = 2.98, 44 deletion carriers). For demographic information see Table 1.

**Table 1.** Participant demographics.

|  | Discovery | Replication | Full |
| --- | --- | --- | --- |
| N | 90 | 92 | 183 |
| Age | 20.71 ± 2.74 years | 20.79 ± 2.98 years | 20.75 ± 2.85 years |
| Sex | 70 female / 19 male / 1 non-binary | 66 female / 26 male | 136 females / 46 males / 1 non-binary |
| ADRA2b Genotype | 43 deletion / 47 non-deletion | 44 deletion / 48 non-deletion | 87 deletion / 95 non-deletion |
| LSAS Score | 30.51 ± 19.40 | 30.69 ± 22.16 | 30.62 ± 20.76 |

### Procedure

Upon arriving in the lab, participants were asked to provide consent and respond to questions regarding vision, hours of sleep, caffeine and alcohol consumption, and history of traumatic brain injury. Additional questionnaires were also administered as part of a standard battery but were not examined for this study. EEG data was subsequently collected while participants performed a change detection task. Following EEG recording, participants performed a stimulus rating task wherein 72 facial stimuli used in the EEG task were rated on scales for valence and arousal. Finally, participants were asked to provide a (∼2ml) saliva sample, collected using an Oragene OG-500 DNA kit (DNA Genotek, Ottawa, ON; http://dnagenotek.com) at the end of each testing session. Genotyping for *ADRA2B* was performed at CAMH in Toronto (1^st^ 125 participants of the final sample) and at the BCCH (BC Children’s Hospital Research Institute) in Vancouver (58 participants of the final sample).

### Stimuli

Seventy-two photographs of 14 actors (12 female) posing angry, neutral, and happy facial expressions were selected from the Karolinska Directed Emotional Faces (KDEF; Lundquist, Flykt, & Ohman, 1998; http://www.facialstimuli.com). These images were converted to grayscale. A Gabor patch was overlaid on each image in a change detection task. Gabor patches were 225 × 305 pixels, with 8 cycles across the vertical dimension. Stimuli were 562 × 762 px (6.244 × 8.467 inches) images that were presented against a gray background on a 24-in monitor with a vertical refresh rate of 60 Hz using Presentation software.

### Change Detection Task

Participants performed a change detection task that was adapted from Wieser, McTeague, and Keil (2012) (Figure 1). *Frequency tagging* was used to flicker overlaid stimuli at opposing frequencies, producing distinct electrocortical signatures, such that power at each driving frequency indexed allocation of cortical resources to that stimulus. In this way we could measure attentional allocation to two distinct stimuli that occupied the same regions of space. Each experimental trial consisted of a face stimulus flickered at one of two frequencies (either 15 or 20 Hz), overlaid by a semi-transparent Gabor patch that flickered at the other frequency (e.g., if the face was flickered at 15 hz, the Gabor patch was flickered at 20hz and vice versa). Both stimuli were presented foveally for 3000 ms. The driving frequencies of face vs. gabor patch were counterbalanced in blocks within subjects across 144 experimental trials. Every trial consisted of 0, 1, or 2 phase reversals of the transparent grating with the first change occurring between 666 ms and 1333 ms and the second change occurring between 1666 ms and 2333 ms. At the end of each trial, a prompt appeared asking the participant to press the number key indicating how many changes were observed (Figure 1b). Whereas the gabor patches were targets, the faces were task-irrelevant. 16 practice trials providing feedback (correct, wrong, or missed) were provided to ensure participants were performing above chance.

**Figure 1.**
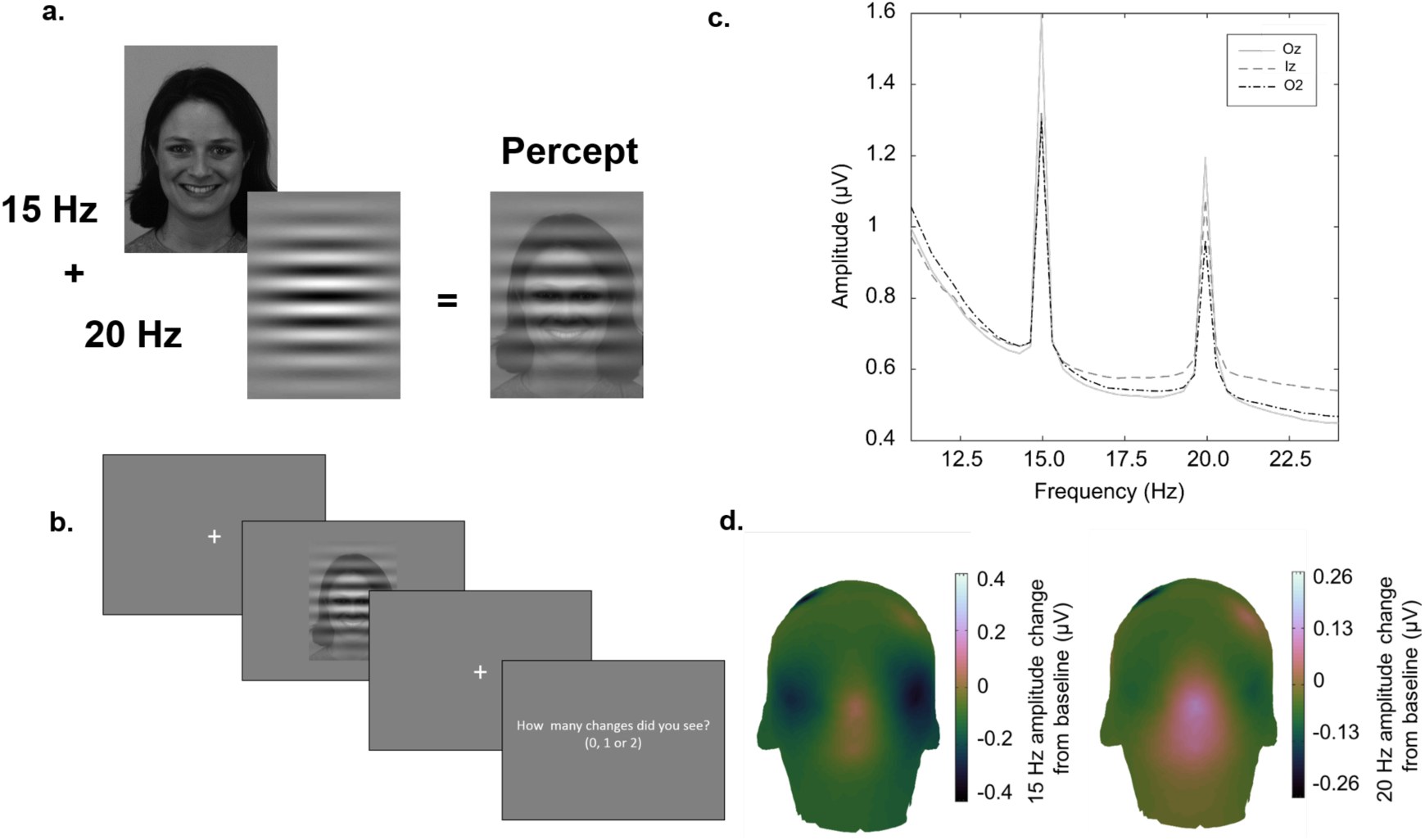
**a.** Stimuli from the change detection task. Target gabor patches and faces showing happy, angry, or neutral expressions were overlaid and flickered at 15 or 20 Hz for three seconds. The frequency of each stimulus type was counterbalanced in blocks within subject. **b**. An example of a trial. For each trial participants had to report whether there were 0, 1 or 2 phase reversals. **c**. Average Fast Fourier Transforms (FFT) for 3 electrodes with the highest signal to noise ratio (SNR) at the peak frequencies. **d**. topographical plot showing 15 and 20 Hz amplitude change from baseline over early visual cortex, computed using the filter-Hilbert method.

### Stimulus rating tasks

Participants viewed the 72 face stimuli once in random order, rating each one of the Self-Assessment Manikin Scale (SAM; Bradley & Lang, 1994) for affective valence and arousal. This self-reporting tool instructs participants to select one of five figures sitting on each of the two 9-point scales to match their experience while viewing a particular stimulus.

Lower scores on the arousal scale indicate a *calm* vs an *exciting* or *agitating* feeling indicated by higher scores. On the valence scale, lower scores more negative ratings while higher scores indicate more positive ratings. Mean ratings were calculated after grouping each into the three emotions (angry, neutral, happy).

### EEG Recording and Analysis

EEG was recorded from 64 active electrodes using the 10-20 Biosemi Active-Two amplifier system. Common mode sense (CMS) and driven right leg (DRL) ground electrodes were placed over medial-parietal cortex. A vertical EOG directly below the right pupil and horizontal EOGs on the left and right outer canthus were attached to record artefactual eye movements, for later correction.

EEGLAB 13.6.5b (Delorme & Makeig, 2004), ERPLAB 6.1 (Lopez-Calderon & Luck, 2014), the PREP pipeline 0.55.2 (Bigdely-Shamlo, Mullen, Kothe, Su, & Robbins, 2015) and MATLAB 2015b (The MathWorks Inc., 2015) were used for offline analysis of data. EEG was predominantly recorded at 256 Hz. EEG was inadvertently recorded at 512 Hz for 4 participants, and 2048 Hz for 2 participants; the data for these participants was downsampled to 256 Hz prior to any pre-processing. The PREP pipeline was used with default parameters to remove line noise and reference all EEG channels to a robust average reference. Bad channels were detected and were replaced with spherical spline interpolated values.

To select electrodes for statistical analysis, signal-to-noise ratio (SNR) was calculated for each electrode as the ratio of the stimulus driving frequencies to the average amplitude of the 20 surrounding frequency bins, excluding those bins immediately adjacent to the driving frequency as well as other frequencies at integer multiples of five to ensure that harmonic frequencies of the driving frequencies were not included in the noise estimate. This SNR calculation follows previous research [e.g., (Liu-Shuang, Norcia, & Rossion, 2014)] with the exception of the additional exclusion of harmonic frequencies. ssVEPs were calculated for each participant as the amplitude at 15 Hz and 20 Hz among the three electrodes with the highest SNR. To obtain 15 Hz and 20 Hz amplitudes for each electrode and trial, the filter-Hilbert method was used with a 12^th^ order Butterworth filter. Obtained amplitudes were then baseline corrected by subtracting the amplitude present at 400 to 200 ms prior to trial onset. As in previous studies (Wieser & Keil, 2011), ssVEP amplitudes for the face and the Gabor patch were then extracted for three time windows (100 to 700 ms; 800 to 1400 ms; 1700 to 2300 ms). For difference score analyses, at each time window a difference score was calculated (*ssVEP*_*Face*_ – *ssVEP*_*Gabor patch*_) such that positive values indicate higher face ssVEP amplitudes than Gabor patch ssVEP amplitudes, and negative values indicate higher Gabor patch ssVEP amplitudes than face ssVEP amplitudes. For the item analysis, this difference score was extracted for each trial, and then averaged across participants to obtain an average measure of attentional allocation for each trial. For the participant-level analysis, the trial-level data was first averaged within each block, and then across blocks, so that each participant has an average ssVEP amplitude for the face, the Gabor patch, and a difference between these two amplitudes, at each specified time window.

### Genotyping

Participants were genotyped for the *ADRA2B* deletion variant. Extraction and genotyping of the DNA was performed at the Neurogenetics Laboratory at the Centre for Addiction and Mental Health in Toronto, Canada for the first 125 participants and at the BC Children’s Hospital Research Institute in Vancouver for the last 58 participants included in the final sample. For each variant, Hardy-Weinberg equilibrium was calculated using an online Hardy-Weinberg Chi Square calculator (Rodriguez, Gaunt, & Day, 2009).

#### Assays

The *ADRA2B* 9bp insertion/deletion variant was assayed using PCR followed by Sanger sequencing. A total of 50ng genomic DNA was combined with 1xAmpliTaq Gold 360 buffer, 2.0 mM Magnesium Chloride, 360GC Enhancer 4ul, 200 uM dNTPs, 0.5 uM Forward primer ACGAAGGTGAAGCGCTTCT and 0.5 uM Reverse primer GGCCAGAAGGAGGGTGTTT, AmpliTaq Gold 360 DNA Polymerse 0.625 U/reaction and total volume 25ul in a 96 well plate. Initial denaturation was at 95°C for 8 min, followed by 38 cycles at 95°C for 50 sec, 60°C for 30 sec and 72°C for 50 sec and a final extension step of 7 min at 72 °C. The PCR products were cleaned up by ExoSAP-IT Express and analyzed by Sanger sequencing (ABI 3130, Applied Biosystems).

## Results

### Genotyping Results

*ADRA2B* genotype frequencies fell within Hardy-Weinberg equilibrium (did not differ from expected frequencies), *X*^*2*^ = 0.03, *p* > .05. Of the 182 individuals, 95 participants did not carry the *ADRA2B* deletion variant, 72 carried one copy of the allele, and 15 were homozygous for the deletion variant. We were unable to obtain reliable genotyping for *ADRA2B* for two participants. For all analyses, based on previous research (de Quervain et al., 2007; Rasch et al., 2009a; Todd et al., 2015; Todd et al., 2013; Todd et al., 2014), homozygote and heterozygote *ADRA2B* deletion carriers were treated as a single group due to the low number of homozygotes.

### Behavioral Results

#### Statistical Analyses

Two-way (*ADRA2B* group) x 3 (emotion) mixed ANOVAs were performed on arousal and valence ratings for face stimuli in the face ratings tasks and for accuracy in the change detection task. Analyses were first conducted independently in the replication and discovery samples. After results were examined in the two separate samples, data from all participants were combined for increased power and analyses were repeated in the full sample. For all ANOVAs reported contrasts were Bonferroni corrected for multiple comparisons and Greenhouse-Geisser corrections were used when sphericity was violated. We report results of analyses of the full sample below. Results of all behavioral analyses for Replication and Discovery samples can be found in Table 2.

**Table 2.** Behavioural Results. Results that are statistically significant at *p* < .05 are in bold, and those that are significant across all samples are bold and underlined.

| | *df* | *F* | *p-value* | $\eta_p^2$ |
| --- | --- | --- | --- | --- |
| **AROUSAL** |  |  |  |  |
| **DISCOVERY SAMPLE** |  |  |  |  |
| **<u>Emotion</u>** | **<u>(1.71, 150.66)</u>** | **<u>45.35</u>** | **<u>0</u>** | **<u>0.34</u>** |
| ADRA2b | (1, 88) | 0.29 | 0.593 | 0 |
| ADRA2b * Emotion | (1.71, 150.66) | 0.16 | 0.818 | 0 |
| **REPLICATION SAMPLE** |  |  |  |  |
| **<u>Emotion</u>** | **<u>(1.74, 154.60)</u>** | **<u>43.14</u>** | **<u>0</u>** | **<u>0.33</u>** |
| ADRA2b | (1, 89) | 0.26 | 0.612 | 0.33 |
| ADRA2b * Emotion | (1.74, 154.60) | 0.07 | 0.916 | 0 |
| **FULL SAMPLE** |  |  |  |  |
| **<u>Emotion</u>** | **<u>(1.74, 311.81)</u>** | **<u>86.57</u>** | **<u>0</u>** | **<u>0.33</u>** |
| ADRA2b | (1, 179) | 0.55 | 0.457 | 0 |
| ADRA2b * Emotion | (1.742, 311.81) | 0.15 | 0.829 | 0 |
| **VALENCE** |  |  |  |  |
| **DISCOVERY SAMPLE** |  |  |  |  |
| **<u>Emotion</u>** | **<u>(1.35, 117.38)</u>** | **<u>387.44</u>** | **<u>0</u>** | **<u>0.82</u>** |
| ADRA2b | (1, 87) | 0.09 | 0.762 | 0 |
| ADRA2b * Emotion | (1.35, 117.38) | 2.33 | 0.12 | 0.03 |
| **REPLICATION SAMPLE** |  |  |  |  |
| **<u>Emotion</u>** | **<u>(1.31, 116.60)</u>** | **<u>554.93</u>** | **<u>0</u>** | **<u>0.86</u>** |
| ADRA2b | (1, 89) | 0.71 | 0.402 | 0.01 |
| ADRA2b * Emotion | (1.31, 116.60) | 0.21 | 0.712 | 0 |
| **FULL SAMPLE** |  |  |  |  |
| **<u>Emotion</u>** | **<u>(1.33, 236.15)</u>** | **<u>899.42</u>** | **<u>0</u>** | **<u>0.83</u>** |
| ADRA2b | (1, 178) | 0.66 | 0.416 | 0 |
| ADRA2b * Emotion | (1.33, 236.15) | 0.63 | 0.471 | 0 |
| **ACCURACY** |  |  |  |  |
| **DISCOVERY SAMPLE** |  |  |  |  |
| Emotion | (2, 176) | 0.9 | 0.41 | 0.01 |
| ADRA2b | (1, 88) | 0.18 | 0.671 | 0 |
| ADRA2b * Emotion | (2, 176) | 2.77 | 0.065 | 0.03 |
| **REPLICATION SAMPLE** |  |  |  |  |
| Emotion | (2, 172) | 2.04 | 0.134 | 0.02 |
| ADRA2b | (1, 86) | 0.07 | 0.788 | 0 |
| ADRA2b * Emotion | (2, 172) | 1.68 | 0.19 | 0.02 |
| **FULL SAMPLE** |  |  |  |  |
| Emotion | (2, 352) | 2.35 | 0.097 | 0.01 |
| ADRA2b | (1, 176) | 0.24 | 0.622 | 0.001 |
| **ADRA2b \* Emotion** | **(2, 352)** | **3.17** | **0.043** | **0.02** |

| **LSAS** |  |  |  |  |
| --- | --- | --- | --- | --- |
| **Discovery sample** |  |  |  |  |
| Emotion | (2, 70) | 2.32 | 0.106 | 0.06 |
| LSAS | (1, 35) | 0.52 | 0.474 | 0.02 |
| LSAS * Emotion | (2,70) | 0.01 | 0.988 | 0 |
| **REPLICATION SAMPLE** |  |  |  |  |
| Emotion | (1.18, 59.03) | 1.72 | 0.195 | 0.03 |
| LSAS | (1, 50) | 1.81 | 0.184 | 0.04 |
| LSAS * Emotion | (1.18, 59.03) | 1.32 | 0.262 | 0.03 |
| **FULL SAMPLE** |  |  |  |  |
| Emotion | (1.29, 111.83) | 3.04 | 0.074 | 0.03 |
| LSAS | (1, 87) | 2.44 | 0.122 | 0.03 |
| LSAS * Emotion | (1.28, 111.83) | 1.27 | 0.272 | 0.01 |

### Arousal Ratings

There was a main effect of emotion, *F*(1.74, 311.81) = 86.57, *p* < .001, 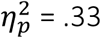, with higher levels of arousal for happy than neutral faces and higher levels of arousal for angry than happy and neutral faces (*p*s < .001). There was no main effect of *ADRA2B, F*(1,179) = 0.55, *p* = .457, 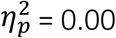 and no emotion x *ADRA2B* interaction, *F*(1.742, 311.81) = 0.15, *p* = .829, 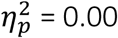. Thus, participants found angry faces to be more arousing than happy faces which in turn were more arousing than neutral faces (Figure 2). There was no evidence that carrying the *ADRA2B* deletion variant influenced subjectively rated arousal of emotional faces.

**Figure 2.**
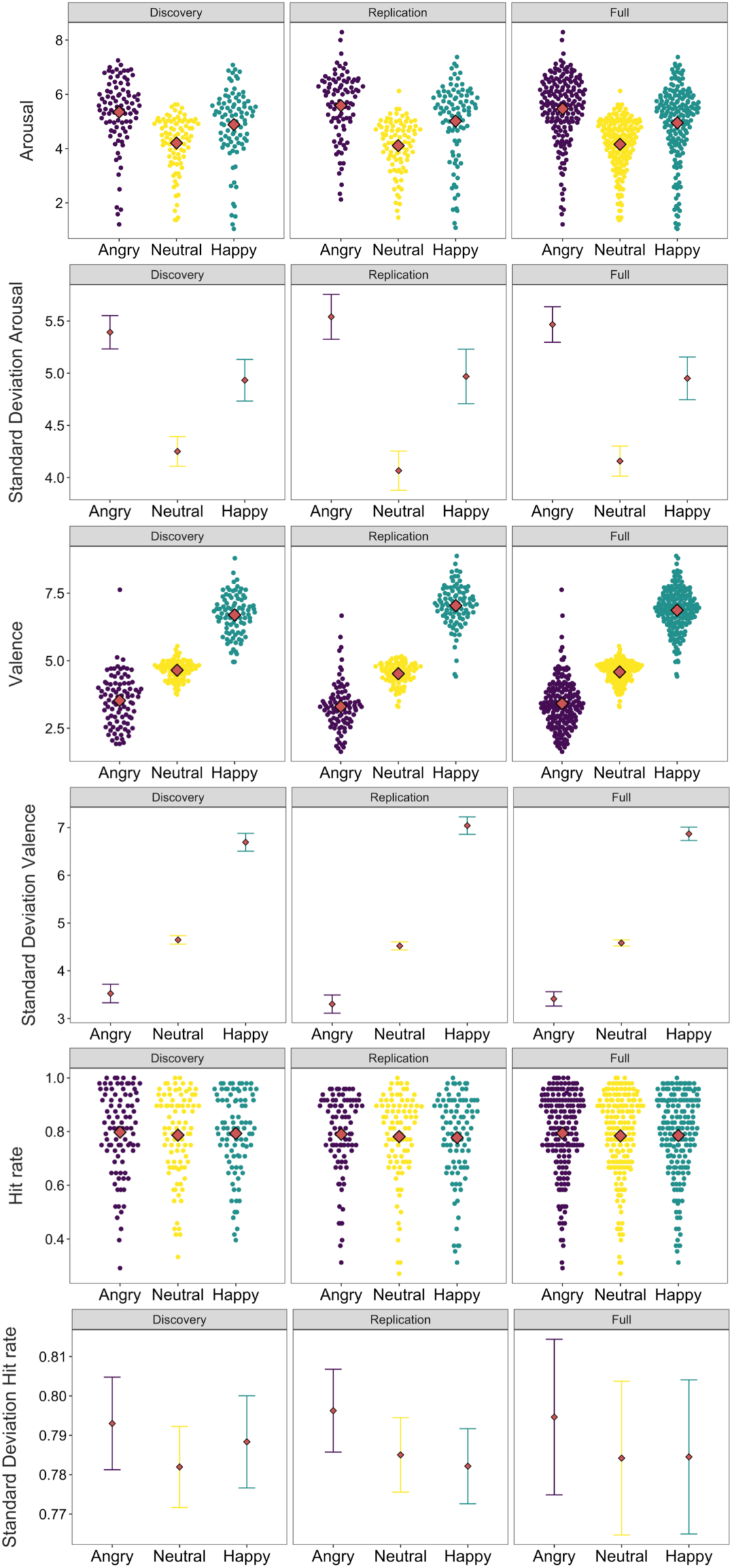
Arousal and valence ratings for face stimuli and accuracy for the change detection task in Discovery, Replication, & Full samles. Angry faces were rated as more arousing than happy faces which were rated as more arousing than neutral faces. Faces were rated as appropriately valenced (lowest numbers are most negative and highest are most positive). And participants did not differ in accuracy between trials with happy, angry or neutral distractors. Lower panels are within-subject error bars.

### Valence Ratings

There was a main effect of emotion, *F*(1.33, 236.15) = 899.42, *p* < .001, 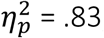, indicating that, as expected, angry faces were rated as negatively valenced and happy faces were rated as positively valenced (*p*’s < .001) There was no main effect of *ADRA2B*, F(1,178) = 0.66, *p* = .416, 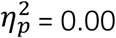 and no emotion x *ADRA2B* interaction, *F*(1.33, 236.15) = 0.63, *p* = .471, 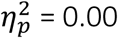. These findings confirm that participants subjectively rate angry, neutral and happy faces as appropriately valenced (Figure 2). There was no evidence that carrying the *ADRA2B* deletion variant influenced subjective experience of stimulus valence.

### Change Detection Accuracy

There was no main effect of emotion, *F*(2,352) = 2.35, *p* = .097, 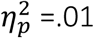, and no main effect of *ADRA2B, F*(1,176) = .24, *p* = .622, 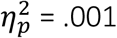 (Figure 2). There was an emotion x *ADRA2B* interaction, *F*(2,352) = 3.17, *p* = .043, 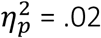, with non-deletion carriers demonstrating greater accuracy in trials with angry face distractors than trials with happy face distractors (paired contrast *p*’s < .05). However, this interaction was non-significant in both the Discovery and Replication samples (see Table 2). Here effect sizes were consistent across samples, indicating the power of the full sample may have been required for a small effect showing facilitation of performance in the presence of angry faces for those who did not carry the deletion variant. This effect may be so small as to be trivial, and would require further replication to merit discussion.

### ssVEP Results

#### Omnibus ANOVA

In order to ascertain the potential presence of competition effects between targets and distractors, we first conducted an omnibus mixed ANOVA on ssVEP power, with *ADRA2B* (deletion carrier vs. non-carrier) as a between-subjects factor and stimulus type (target or distractor), emotion (angry, neutral, happy) and time window (early, middle, late) as within subject factors. All results of this ANOVA are reported in Table 3. There was a main effect of stimulus type, *F*(1,180) = 14.34, *p* < .001, 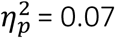 such that there were higher amplitudes in response to the Gabor targets than to the face distractors, indicating biased competition favoring target stimuli. There was a consistently observed stimulus type x emotion interaction, indicating the presence of affectively-biased competition, *F*(2,360) = 13.06, *p* < .001,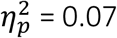. Paired contrasts revealed that whereas amplitudes were greater for targets in the presence of all distractor types (*p*’s < .05), the differences were smaller in trials with happy face distractors. There was also a time window x stimulus type interaction, *F*(1.21,218.69) = 14.75, *p* = .000, 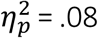. Pairwise contrasts indicated greater power differences between targets and face distractors in window 1 and 3 than in window 2 (*p*s < .005). Importantly, there was no main effect of *ADRA2B, F*(1, 180) = .88, *p* = .350, 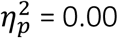 and no hypothesized stimulus type x emotion x *ADRA2B* interaction, *F*(2, 360) = 1.49, *p* = .227, 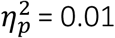.

**Table 3.**
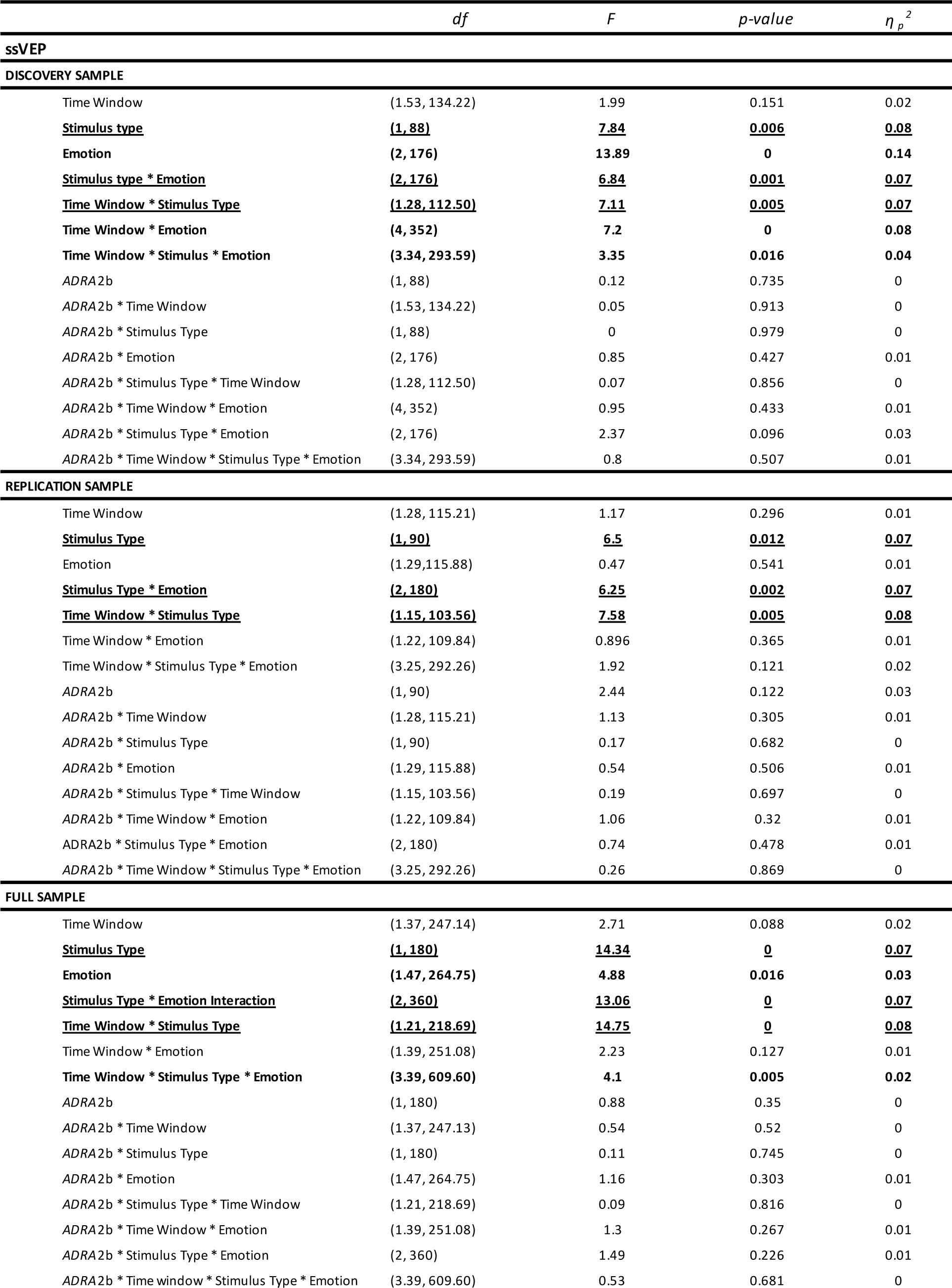
ssVEP results. Results that are statistically significant are in bold, and those that are significant across all samples are bold and underlined.

In summary, in all samples there was a pattern of higher amplitudes for target stimuli, indicating competition effects indicating participants maintained greater attention to task-relevant stimuli than emotionally salient distractors. We also reliably observed an interaction between stimulus type and distractor expression, indicating that the difference in power between targets and distractors was reduced in the presence of happy faces. We further observed that the differences in ssVEP power varied by time window. Notably, there was no evidence of any effect of *ADRA2B* genotype.

#### Difference Scores

To follow up on the interaction between facial expression and stimulus type, and thus better probe the characterize effects of emotion on competition for neurocognitive resources between targets and distractors, we next calculated a difference score between targets and distractors for each emotion expression in each time window. Thus, larger difference scores would indicate greater competition between targets and distractors and vice versa. Moreover, while our previous studies have found robust effects of *ADRA2B* on emotion/cognition interactions in samples of this size or smaller, suggesting our study was not underpowered to find equivalent effects, we wished to further probe the reliability of the null effects observed for *ADRA2B.* We thus next conducted a Bayesian ANOVA on the full sample using JASP (JASP Team, 2018) with default priors. One participant who showed difference scores - 6.5 standard deviations from the mean was excluded from these analyses. The pattern of results did not differ when the outlier was included.

First, we conducted a mixed ANOVA with *ADRA2B* (deletion carrier vs. non-carrier) as a between subject factor and emotion (angry, neutral, happy) and time window (early, middle, late) as within subject factors. Consistent with the time window by stimulus type interaction observed in the previous analysis, there was a main effect of time window, *F*(1.53, 274.06) = 12.55, *p* = .000, η_*p*_^2^ = .07 such that there were overall largest difference scores in the third time window than in either of the first 2 (*p*s < .001). There was also a main effect of emotion, *F*(2, 358) = 7.59, *p* = .001, η_*p*_^2^= .04. Contrasts revealed that difference scores were smaller in trials with happy faces, indicating greater competition for neurocognitive resources from happy faces than from neutral (p < .001) or, marginally, angry (*p* = .06) faces, which in turn did not differ from each other (*p* > 1) (Figure 3). In this analysis this finding was qualified by a time window x emotion interaction, *F*(3.76, 672.63) = 3.30, *p* = .012, η_*p*_^2^ = .02, indicating that the greatest happy face bias effect was observed in time window 2 (*p*s < .05). Crucially, there was no main effect of *ADRA2B, F*(1, 179) = .08, *p* = .774, η_*p*_^2^= .00, no emotion x *ADRA2B* interaction, *F*(2, 358) = .728, *p*= .483, η_*p*_^2^ = .00 (Figure 3), nor was there a time window x emotion x *ADRA2B* interaction, *F*(3.76, 672.63) = .34, *p* = .850, η_*p*_^2^ = .00. As a subsequent item analysis (described below) revealed that two happy face stimuli elicited ssVEP difference scores that were greater than 2.5 SD from the mean, we next reran the analysis with the trials using those faces removed. The pattern of results did not change with the happy face outliers removed. Crucially, the main effect of emotion showing greater competition for happy faces (*p* = .010, η_*p*_^2^ = 0.02) and the emotion by time window interaction (*p* = .006, η_*p*_^2^ = 0.02), indicating this effect was most pronounced in the middle time window, were still observed. Thus, the happy face distraction effect was not driven by outliers.

**Figure 3.**
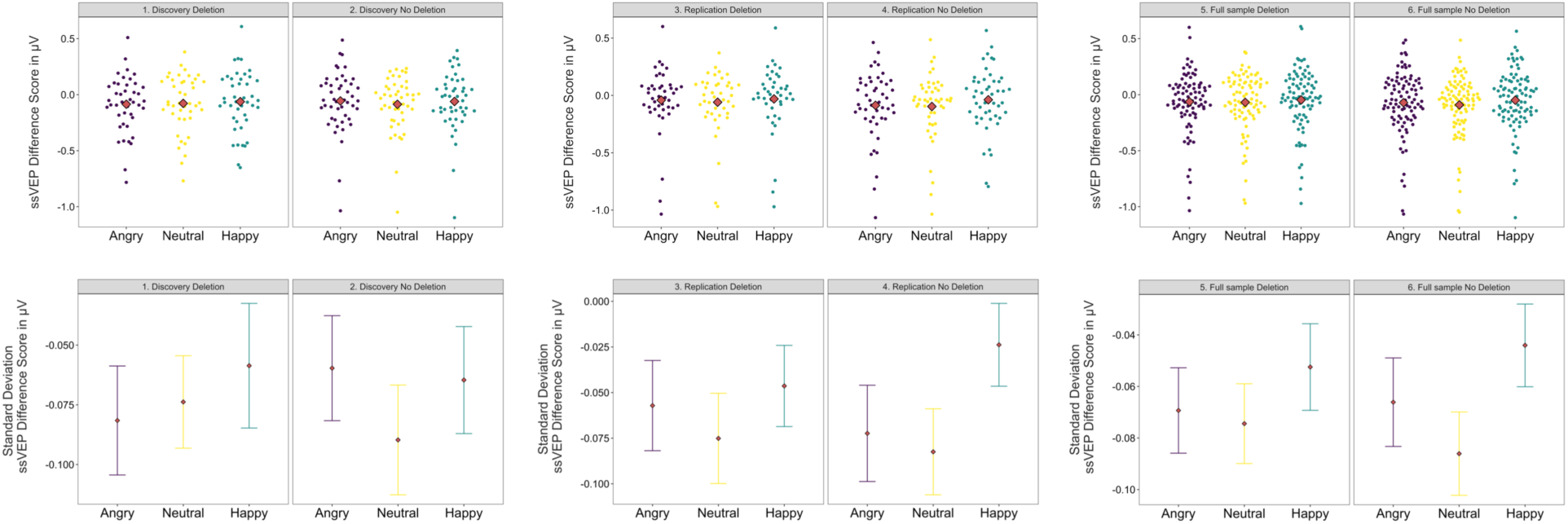
ssVEP difference scores (*ssVEP*_*Face*_− *ssVEP*_*Gabor patch*_) for trials with distractors for each facial emotion for the Discovery, Replication, and Full samples by *ADRA2B* group (deletion variant carriers and non-carriers). Overall more cortical resources were allocated to task relevant stimuli than emotional face distractors. There were reduced difference scores in trials with happy faces, indicating greater competition from happy faces. There was no influence of carrying the *ADRA2B* deletion variant on this pattern of results.

Analysis of the effects of a Bayesian Mixed ANOVA across matched models revealed a main effect of time window (*BF*_*Inclusion*_= 15302.41), a main effect of face emotion (*BF*_*Inclusion*_= 1064.12) and, in this analysis, no interaction between these two factors (*BF*_*Inclusion*_= 0.003). There is some evidence against the main effect of *ADRA2B* (*BF*_*Inclusion*_= 0.31) and its interaction with face emotion (*BF*_*Inclusion*_= 0.27) and its interaction with time window (*BF*_*Inclusion*_= 0.02). There is considerable evidence against a three-way interaction among these factors (*B*⟁_*Inclusion*_= 0.002). These results demonstrate that there is evidence against ADRA2B modulating the degree to which emotional stimuli influence biased competition.

In summary, effects that were consistent across all frequentist and Bayesian analyses of ssVEP amplitude indicated that, within each trial, competition for neurocognitive resources favouring target stimuli over face distractors increased over time. Competition was also modulated by the facial expression of the distractor, such that smiling faces captured more neurocognitive resources from targets than angry or neutral faces. Effects of time window on emotionally-biased competition were not reliable. Importantly, there was no evidence for – and some positive evidence against — our hypothesis that putative greater NE availability resulting from carrying the ADRA2B deletion variant influenced emotionally-biased competition between targets and distractors.

#### Effects of Social Anxiety

While reliable, the observed effects of emotional valence on difference scores were small. In order to examine whether the higher levels of competition for neurocognitive resources was modulated by social anxiety, as has been previously observed (Wieser et al., 2012), we next probed effects of emotion on the difference scores in the 25% of our sample that was highest (n = 45), and the 25% that was lowest (n =44) in social anxiety based on LSAS scores. Mean scores for social anxiety (LSAS) were (*M* = 9.68, *SD* = 7.39) in the low anxiety group and (*M* = 58.87, *SD* = 14.20) in the high anxiety group.

A 2×3×3 (LSAS group x Emotion x Time window) mixed ANOVA was performed on ssVEP difference scores described above. As previously reported, there was no main effect of emotion, *F*(1.29, 112.56) = 3.26, *p* = .063,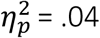. There was a main effect of time window, F(1.60, 139.57) = 5.84, p = .007, 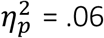 but no time window x emotion interaction F(4, 348) = 2.13, p =.076,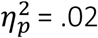. Finally, there was no main effect of LSAS, *F*(1, 87) = 2.44, *p* = .122, 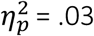 and no emotion x LSAS interaction, *F*(1.29, 112.56) = 1.06, *p* = .322, 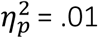.

To confirm the null effect of social anxiety, analysis of the effects of a Bayesian Mixed ANOVA across matched models again revealed a main effect of time window (*BF*_*Inclusion*_= 3.45), a main effect of face emotion (*BF*_*Inclusion*_= 71.60). There was inconclusive evidence for both a main effect of social anxiety and an interaction between social anxiety and face emotion (*BF*_*Inclusion*_= 0.86; *BF*_*Inclusion*_= 0.418, respectively). All remaining interactions revealed null effects (all *BF*_*Inclusion*_< 0.07). Thus there was no evidence suggesting that social anxiety influenced affectively-biased competition in this sample of participants.

#### Trial by trial examination of low-level feature effects, valence and arousal

An outstanding question concerned whether the effects of emotional valence on biased competition were due to the emotional content of the stimuli or capture of attention by low level features, such as contrast, that might differentiate happy faces from neutral and angry faces. To probe this question, we extracted a variety of image metrics. First, we used the Image Processing Toolbox packaged with MATLAB 8.6.0 (The MathWorks, Natick, MA) to calculate whole image contrast, average local image contrast within distinct square regions of different sizes (8×8, 16×16, 32×32, 64×64, and 128×128 regions), edges, and average log luminance. Second, we used the SHINE toolbox (Willenbockel et al., 2010) to calculate low and high spatial frequency content. Third, we used the Visual Saliency Toolbox (Walther & Koch, 2006) to calculate visual salience and the Euclidean distance between the first and second predicted fixations. Each of these measures were correlated with ssVEP difference score values. Contrast was calculated by finding the standard deviation of the grayscale pixel values. Edges was calculated with MATLAB’s edge function, using a Canny filter with a threshold of 0.5. Average log luminance was calculated as the exponential of the mean value of the logarithm of the grayscale pixel values. Spatial frequency content was calculated using the sfplot function from the SHINE toolbox, which extracts the energy at each spatial frequency in the image. From the output of this function, low frequency content was calculated as the mean energy of frequencies below the median frequency, and high frequency content was calculated as the mean energy for the remaining frequencies. A single image was removed from the analysis because it was identified visually as an outlier and had a face-Gabor ssVEP difference greater than 2.5 standard deviations from the mean. Results showed that only edges may have marginally influenced the face-Gabor ssVEP differences (r = 0.18, p = .131, 95% CI [- 0.05, 0.40]), while all other image properties very clearly did not correlate with this measure (*p*’s > .270) (Supplementary Figure 1). For each trial we also extracted subjective valence and arousal ratings averaged across participants.

Next, we conducted a one-way ANOVA to test whether the number of edges differed between facial emotion categories. If the ANOVA revealed that there was no difference among the face emotions on the amount of edges in the images, then our primary result demonstrating modulation of amplitude differences by face emotion should not be due to differences in the amount of edges in the image. However, the ANOVA revealed that facial emotion category had a significant impact on the number edges in an image, *F*(2, 68) = 4.80, *p* = .011, 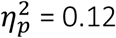). To determine whether this difference in number of edges across emotion categories could be responsible for the observed Face-Gabor patch ssVEP differences, above and beyond the influence of emotional arousal and valence, we performed a regression with edges, as well as ratings of arousal, valence, and the interaction of arousal and valence for each face stimulus as predictors of Face-Gabor patch ssVEP difference. The full regression model showed no significant effect of edges, β = 0.13, *t*(66) = 1.12, *p* = .265, and did not differ from a more parsimonious model with only arousal, valence, and their interaction as predictors, F(1,66) = 1.26, *p* = .265, so we will describe the results of the more parsimonious model. Face-Gabor ssVEP difference was significantly predicted by valence, β = 0.47, *t*(67) = 3.03, *p* = .003, but not arousal, β = 0.04, *t*(67) = 0.27, *p* = .791. There was a trend level interaction, β = −0.45, *t*(67) = −1.87, *p* = .065, which we illustrate in Figure 4 via simple slopes of the regression of Face-Gabor ssVEP difference on arousal at three different levels of valence (Mean valence, 1 SD below mean valence, and 1 SD above mean valence.) Follow-up tests of these three simple slopes using bootstrapping with 9999 bootstrap replicates and calculating the BCa confidence interval (DiCiccio & Efron, 1996) reveal that arousal predicts higher Face-Gabor ssVEP difference only among low valence face stimuli (i.e. negative faces), 95% CI [0.12, 0.79]. Face-Gabor patch difference is not significantly predicted by arousal at mean valence (95% CI [-0.25, 0.31]), or high valence, (95% CI [-1.05, 0.15]). In sum, though edges differed across stimulus emotion categories, it did not significantly predict Face-Gabor patch ssVEP differences. Instead, we found that once again valence predicted difference scores, confirming the happy face advantage. In addition, higher stimulus arousal results in higher Face-Gabor ssVEP differences (i.e. a relatively higher ssVEP in response to face stimuli), but only among more negatively valenced stimuli. Thus neither any of the low level features we measured nor subjective ratings of arousal accounted for the happy face advantage.

**Figure 4.**
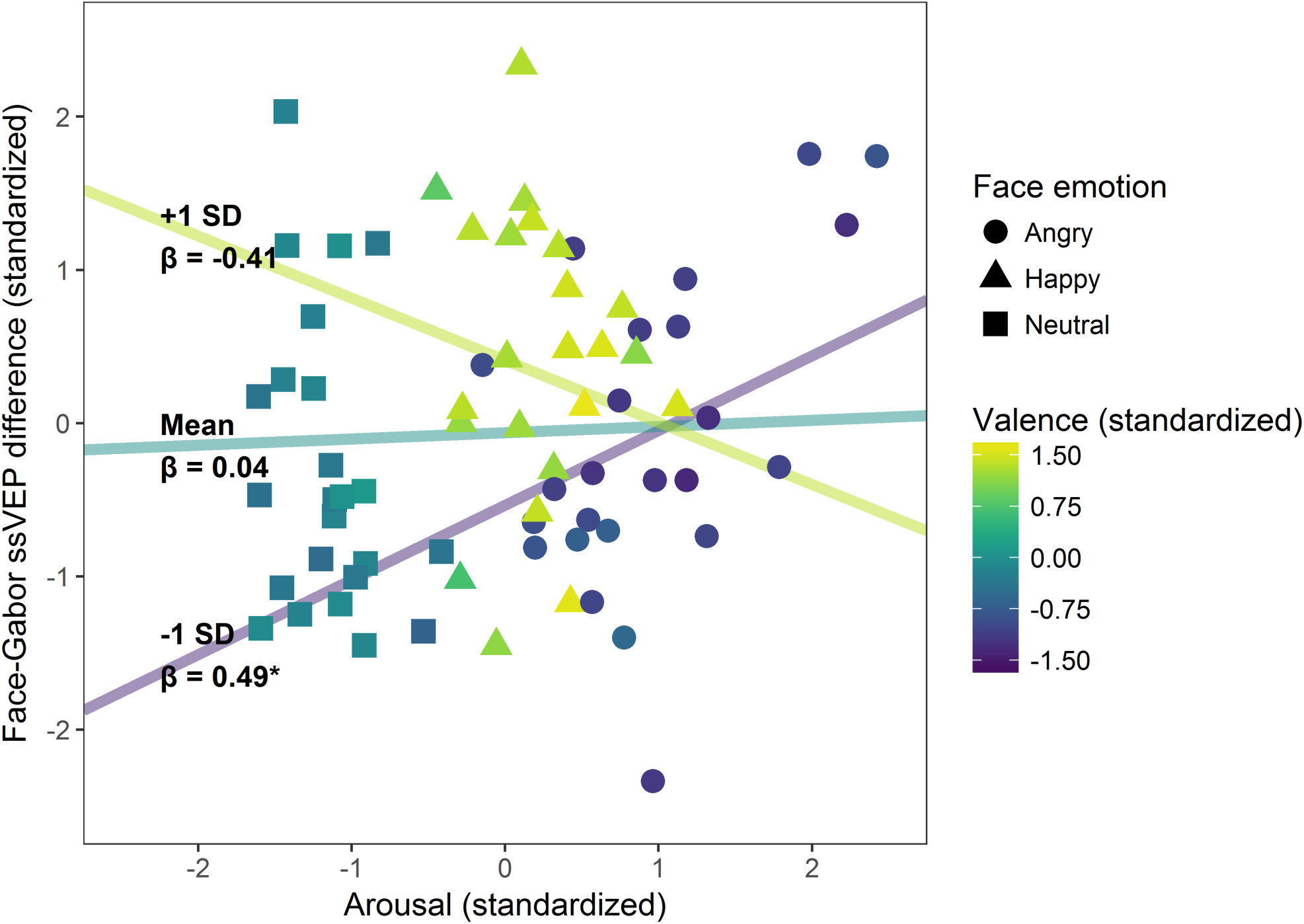
Regression of Face-Gabor ssVEP differences on stimulus arousal, valence, and the interaction between arousal and valence. A follow-up on the trend-level interaction between arousal and valence demonstrated a significant positive relationship between arousal and ssVEP differences, but only among more negatively valenced stimuli (i.e. angry faces). This suggests that participants devote more attentional resources to highly arousing (task-irrelevant) stimuli when they are negative, but that arousal has no influence on attention when viewing neutral or positive task-irrelevant stimuli.

## Discussion

In the present study, we used ssVEPs to examine effects of a deletion variant in *ADRA2B*, a common genetic variation thought to influence norepinephrine availability, on affectively-biased competition in the visual cortex. We further examined within-subject effects of emotional valence in a large sample of healthy young adults. Counter to our hypotheses, carrying a deletion variant of the *ADRA2B* gene had no influence on competition for population-level neuronal responses between task-relevant stimuli and affectively salient distractors. Thus, the evidence suggests that naturally occurring variation in NE levels putatively associated with this genetic variation neither drive nor modulate such affectively-based visual cortex competition when attention must be sustained over seconds. In the group as a whole, the presence of happy faces captured cortical resources from target stimuli, particularly at middle latencies, indicating biased competition for smiling faces at the population level of neuronal competition. Although photographs of happy faces did differ from those of angry and neutral faces in the number of edges they contained, follow-up analyses showed that the effect of emotional expression on biased competition between happy faces and other facial expressions was not due to differences in any of the low-level features we measured or in subjective ratings of arousal.

Our own previous research has found robust effects for the *ADRA2B* deletion variant influencing affective biases in attention and memory. Building on findings that carrying the deletion variant influences emotional enhancement of memory (de Quervain et al., 2007), we have found that carrying this variation enhances rapid attentional selection for emotionally relevant words, enhances the link between subjective emotional arousal in encoding and memory accuracy and confidence, and enhances the subjective perception of the relative vividness (signal to noise ratio) of emotionally salient images (Todd et al., 2015; Todd et al., 2013; Todd et al., 2014). It has also been associated with differences in responsiveness of the amygdala and VMPFC to enhanced processing of salient stimuli (Rasch et al., 2009b; Todd et al., 2015; Todd et al., 2013; Todd et al., 2014), which can drive affectively enhanced processing in the visual cortex.

The present study was motivated by the hypothesis that putatively higher levels of NE availability in deletion carriers would facilitate biased visual cortex competition favoring affectively salient stimuli, providing some mechanistic understanding of our previously-observed results. This was not the case. It is worth noting that all of our previous studies on the influence of *ADRA2B* on attention and subjective perception tapped relatively rapid processes, whereas the effects of biased competition measured here emerged more slowly. Moreover, previous studies measuring effects of the deletion variant on emotional attention and perception engaged *subjective judgments* of emotional content (Ehlers, Ross, & Todd, 2018), or the relative level of noise overlaid on a visual stimulus (Todd et al., 2015), or rapid engagement of selective attentional prioritization of emotional content (Todd et al., 2013). In contrast, the present study required sustained attention to a neutral stimulus over the course of seconds, and therefore continuous inhibition of distraction by the emotional expression. The LC/NE system is thought to influence emotional effects on perception, attention and memory via amygdala modulation of other brain regions [for review see (Ehlers & Todd, 2017; Markovic et al., 2014; Roozendaal, Luyten, de Voogd, & Hermans, 2016)]. Given electrophysiological evidence that amygdala neurons are phasically activated – showing transient increase in firing at the onset of salient or novel stimuli stimuli (E. J. Kim et al., 2018)— it may be that the sustained pulse of emotional distractors did not elicit amygdala activation that would differentiate deletion carriers and non-carriers. If true, one boundary condition of effects of *ADRA2B* on subjective perception and attention would it does not influence sustained or ongoing processes. This hypothesis should be tested directly in future research.

It is also notable that our previous research found that, in contrast to findings of emotionally enhanced activation in the ventromedial prefrontal cortex and the amygdala (Rasch et al., 2009b; Todd et al., 2015), deletion carriers did not differ from non-carriers in patterns of enhanced visual cortex activation for emotionally salient stimuli. These findings are convergently consistent with the conclusion that effects of NE on emotionally-biased processing do not function via visual cortex activity.

The results of a Bayesian ANOVA further suggested our null finding was conclusive, despite the relatively small sample size for a targeted polymorphism study. Our power analysis indicated that the study was amply powered based on the weakest effect size of our previous studies examining *ADRA2B.* Thus, we conclude that commonly observed differences related to NE activity do not play a role in modulating biased competition in the visual cortex, at least not for sustained firing patterns measured here. However, it is worth noting that previous studies investigating effects of *ADRA2B* have employed more arousing affective stimuli than emotionally expressive faces, using images and words that evoke more physiological arousal [e.g., (A. K. Anderson, Yamaguchi, Grabski, & Lacka, 2006). Thus, it is possible that a higher level of stimulus-evoked arousal may be required for differences in NE functioning to influence biased competition effects.

We did, however, find reliable evidence that the smiling faces we used captured more visual cortex resources, indexing greater attention, from task-relevant stimuli than angry or neutral faces. These findings are inconsistent with a prevalent “standard hypothesis” in affective science, which has proposed that threatening stimuli such as angry faces are universally prioritized because of an evolutionary advantage for rapid threat detection (Ohman, Flykt, & Esteves, 2001; Ohman, Lundqvist, & Esteves, 2001). However, findings of attentional prioritization of threatening relative to rewarding facial expressions have been repeatedly challenged on several fronts. First, there is an extensive literature documenting systematic individual differences in the direction of biases [for review see (Todd, Cunningham, Anderson, & Thompson, 2012)]. Valence preferences can also vary with developmental stage, such that young children show far more sensitivity and prioritization of happy faces than young adults (Picardo, Baron, Anderson, & Todd, 2016; Todd, Evans, Morris, Lewis, & Taylor, 2011). In the present study, this pattern did not reflect the individual differences in *ADRA2B* genotype and social anxiety we predicted and measured in this sample. This may be because previous studies finding social anxiety effects used participants with less extreme levels of social anxiety and less variability in the high social anxiety participants (Wieser et al., 2012). However, it is likely that given the degree of variability observed (Figure 2), and the modest effect size, this pattern is subject to individual differences linked to other traits.

Second, ERP studies of emotional effects on visual processing have reported different early patterns of response and time courses for positive vs. negative stimuli, potentially reflecting differences in contributions of reward circuitry to valenced responses. There is evidence that the C1 component, which is generated by striate cortex as rapidly as within 50 ms of stimulus onset, responds preferentially to negatively valenced over neutral stimuli in a study where positive valence was not tested (West, Anderson, & Pratt, 2009), and to monetary loss relative to monetary gain (Rossi et al., 2017). In contrast, later time windows have shown enhanced activation for both monetary gain and loss over neutral stimuli. Other studies have also found reward-related responses, putatively mediated by estimated anterior cingulate activity, between 200 and 300 ms following stimulus onset (Hickey, Chelazzi, & Theeuwes, 2010). Thus, whereas very rapid visual cortex activity appears to favor negatively valenced stimuli, activity that may reflect more sustained re-entrant processing may favor salient stimuli in general. Our findings showing that the greater competition for happy faces was greatest in the latest time window, allowing for potential feedback from reward circuitry, are not inconsistent with these findings.

Third, studies looking at overall within-subject effects of valenced stimuli on attention have yielded conflicting results. After controlling for low-level features, some researchers have observed consistent happiness superiority effects as we did (Becker, Anderson, Mortensen, Neufeld, & Neel, 2011). One suggestion has been that, after controlling for low-level perceptual features, it is arousal rather than valence that drives attentional prioritization (Juth, Lundqvist, Karlsson, & Ohman, 2005). However, our present findings are not consistent with this interpretation. In the present study, participants rated angry faces as more arousing than happy faces, and arousal accounted for ssVEP difference scores in angry faces only — there was no relationship between arousal and biased competition favoring happy faces. Previous findings of happy face prioritization have been interpreted as reflecting easier visual discrimination of the unambiguous communication that is afforded by smiling faces (Becker et al., 2011). This interpretation is the most consistent with our results, where rewarding content that was unrelated to arousal captured attention.

In summary, our findings are consistent with robust evidence that in some contexts happy facial expressions, which are experienced as rewarding, are prioritized over angry ones. These findings did not reflect low-level features nor did they reflect patterns of subjectively rated arousal. Nonetheless, such valence effects are subject to numerous boundary conditions that constrain our interpretation. We measured a wide range of low level features, including contrast at a multiple spatial scales, edges, luminance, and a measure of visual saliency that predicts eye movement behaviour, and predicted eye-movement behaviour, and none of these predicted ssVEP difference scores; however, in schematic faces the happy face advantage has been linked to more subtle featural confounds (Purcell & Stewart, 2010), such as the relative orientation of face and mouth contours (Stein & Sterzer, 2012). In the case of photographic faces, prioritization of angry vs happy faces can vary depending on the stimulus set used (Savage, Becker, & Lipp, 2016; Savage, Lipp, Craig, Becker, & Horstmann, 2013). Future research will be required to establish whether the increased biased competition from happy faces we observed generalizes to other affective stimulus sets.

Finally, although examination of social anxiety was not the focus of this study, we failed to replicate previous findings that social anxiety modulated the biased competition effect in a larger sample (Wieser et al., 2012). Unlike the previous study, we did not screen for high and low anxiety, and our high-anxiety group had lower levels of mean anxiety than that of the previous study. Thus, it may be effects would be replicated in samples where anxiety levels are more extreme.

In sum, our findings provide compelling evidence that affectively biased competition in sustained attention is not modulated by individual differences in noradrenergic activity related to alpha2b receptor function. Moreover, although we observed consistent affectively-biased competition for smiling faces, the present findings add to growing evidence indicating multiple boundary conditions for previously observed valence effects. Studies using larger sample sizes for within-subject designs, investigation of individual differences, and increased use of tools such as Bayesian analyses to probe the conclusiveness of null results – along with increased willingness to publish null results – will play an important role refining our understanding of cognition-emotion interactions.

## Supporting information

Supplemental correlation plots

## Acknowledgments

This research was supported by Canadian Institutes for Health Research (CIHR) Operating Grant #491746, National Engineering and Research Council (NSERC) Discovery Grant RGPIN-2014-04202, and Canadian Foundation for Innovation Leaders Opportunity Fund grant F13-03917. Individual researchers were supported by a NSERC doctoral fellowship to K. H. Roberts and CIHR New Investigator Salary awards and Michael Smith Foundation for Health Research Scholar awards to both C.J.D. Ross and R.M. Todd. We would like to thank Drs. Todd Handy and Julia Kam as well as Mana Ehlers and Lena Fürst for their generous contributions to this project.

## Data accessibility statement

All data and analysis scripts are available on Open Science Framework: https://osf.io/n3g6e/?view_only=458920fd64c048028c651f78f7192f55

## References

Anderson, A. K., Yamaguchi, Y., Grabski, W., & Lacka, D. (2006). Emotional memories are not all created equal: evidence for selective memory enhancement. Learn Mem, 13(6), 711–718. doi:lm.388906[pii]10.1101/lm.388906

Anderson, B. A. (2016). What is abnormal about addiction-related attentional biases? Drug and alcohol dependence, 167, 8–14. doi:10.1016/j.drugalcdep.2016.08.002

Bar-Haim, Y., Lamy, D., Pergamin, L., Bakermans-Kranenburg, M. J., & van, I. M. H. (2007). Threat-related attentional bias in anxious and nonanxious individuals: a meta-analytic study. Psychological Bulletin, 133(1), 1–24. doi:10.1037/0033-2909.133.1.1

Becker, D. V., Anderson, U. S., Mortensen, C. R., Neufeld, S. L., & Neel, R. (2011). The face in the crowd effect unconfounded: happy faces, not angry faces, are more efficiently detected in single-and multiple-target visual search tasks. J Exp Psychol Gen, 140(4), 637–659. doi:10.1037/a0024060

Berridge, C. W., & Waterhouse, B. D. (2003). The locus coeruleus-noradrenergic system: modulation of behavioral state and state-dependent cognitive processes. Brain research. Brain research reviews, 42(1), 33–84.

Carretie, L. (2014). Exogenous (automatic) attention to emotional stimuli: a review. Cogn Affect Behav Neurosci, 14(4), 1228–1258. doi:10.3758/s13415-014-0270-2

Dalgleish, T., Taghavi, R., Neshat-Doost, H., Moradi, A., Canterbury, R., & Yule, W. (2003). Patterns of processing bias for emotional information across clinical disorders: a comparison of attention, memory, and prospective cognition in children and adolescents with depression, generalized anxiety, and posttraumatic stress disorder. Journal of clinical child and adolescent psychology: the official journal for the Society of Clinical Child and Adolescent Psychology, American Psychological Association, Division 53, 32(1), 10–21. doi:10.1207/S15374424JCCP3201_02

de Quervain, D. J., Kolassa, I. T., Ertl, V., Onyut, P. L., Neuner, F., Elbert, T., & Papassotiropoulos, A. (2007). A deletion variant of the alpha2b-adrenoceptor is related to emotional memory in Europeans and Africans. Nat Neurosci, 10(9), 1137–1139. doi:nn1945[pii]10.1038/nn1945

de Quervain, D. J., & Papassotiropoulos, A. (2006). Identification of a genetic cluster influencing memory performance and hippocampal activity in humans. Proc Natl Acad Sci U S A, 103(11), 4270–4274. doi:0510212103[pii]10.1073/pnas.0510212103

Derryberry, D., & Reed, M. A. (1994). Temperament and attention: orienting toward and away from positive and negative signals. Journal of Personality and Social Psychology, 66(6), 1128–1139.

Ehlers, M. R., Ross, C. J. D., & Todd, R. M. (2018). The influence of the noradrenergic/stress system on perceptual biases for reward. Cogn Affect Behav Neurosci. doi:10.3758/s13415-018-00657-0

Ehlers, M. R., & Todd, R. M. (2017). Genesis and maintenance of attentional biases: The role of the Locus Coeruleus-Noradrenaline system. Neural Plasticity, 1-15.doi:https://doi.org/10.1155/2017/6817349

Ehlers, M. R., & Todd, R. M. (2018). Adaptation and noradrenergic genetic variations influence emotional categorization. Paper presented at the 2018 International Conference on Learning and Memory.

Fairfield, B., Mammarella, N., Fontanella, L., Sarra, A., D’Aurora, M., Stuppia, L., & Gatta, V. (2019). Aging and the Combined effects of ADRA2B and CB1 deletions on Affective Working Memory. Sci Rep, 9(1), 4081. doi:10.1038/s41598-019-40108-5

Gelbard-Sagiv, H., Magidov, E., Sharon, H., Hendler, T., & Nir, Y. (2018). Noradrenaline Modulates Visual Perception and Late Visually Evoked Activity. Curr Biol, 28(14), 2239–2249 e2236. doi:10.1016/j.cub.2018.05.051

Grabitz, C. R., Button, K. S., Munafo, M. R., Newbury, D. F., Pernet, C. R., Thompson, P. A., & Bishop, D. V. M. (2018). Logical and Methodological Issues Affecting Genetic Studies of Humans Reported in Top Neuroscience Journals. J Cogn Neurosci, 30(1), 25–41. doi:10.1162/jocn_a_01192

Hedge, C., Powell, G., & Sumner, P. (2018). The reliability paradox: Why robust cognitive tasks do not produce reliable individual differences. Behav Res Methods, 50(3), 1166–1186. doi:10.3758/s13428-017-0935-1

Hickey, C., Chelazzi, L., & Theeuwes, J. (2010). Reward changes salience in human vision via the anterior cingulate. J Neurosci, 30(33), 11096–11103. doi:10.1523/JNEUROSCI.1026-10.2010

Juth, P., Lundqvist, D., Karlsson, A., & Ohman, A. (2005). Looking for foes and friends: perceptual and emotional factors when finding a face in the crowd. Emotion, 5(4), 379–395. doi:10.1037/1528-3542.5.4.379

Kastner-Dorn, A. K., Andreatta, M., Pauli, P., & Wieser, M. J. (2018). Hypervigilance during anxiety and selective attention during fear: Using steady-state visual evoked potentials (ssVEPs) to disentangle attention mechanisms during predictable and unpredictable threat. Cortex; a journal devoted to the study of the nervous system and behavior, 106, 120–131. doi:10.1016/j.cortex.2018.05.008

Kim, E. J., Kong, M. S., Park, S. G., Mizumori, S. J. Y., Cho, J., & Kim, J. J. (2018). Dynamic coding of predatory information between the prelimbic cortex and lateral amygdala in foraging rats. Sci Adv, 4(4), eaar7328. doi:10.1126/sciadv.aar7328

Kim, Y. J., Grabowecky, M., Paller, K. A., Muthu, K., & Suzuki, S. (2007). Attention induces synchronization-based response gain in steady-state visual evoked potentials. Nat Neurosci, 10(1), 117–125. doi:nn1821[pii]10.1038/nn1821

Liu-Shuang, J., Norcia, A. M., & Rossion, B. (2014). An objective index of individual face discrimination in the right occipito-temporal cortex by means of fast periodic oddball stimulation. Neuropsychologia, 52, 57–72. doi:10.1016/j.neuropsychologia.2013.10.022

Makaritsis, K. P., Johns, C., Gavras, I., & Gavras, H. (2000). Role of alpha(2)-adrenergic receptor subtypes in the acute hypertensive response to hypertonic saline infusion in anephric mice. Hypertension, 35(2), 609–613.

Markovic, J., Anderson, A. K., & Todd, R. M. (2014). Tuning to the significant: Neural and genetic processes underlying affective enhancement of visual perception and memory. Behavioural brain research, 259, 229–241. doi:10.1016/j.bbr.2013.11.018

Mather, M., Clewett, D., Sakaki, M., & Harley, C. W. (2016). Norepinephrine ignites local hotspots of neuronal excitation: How arousal amplifies selectivity in perception and memory. Behav Brain Sci, 39, e200. doi:10.1017/S0140525X15000667

McTeague, L. M., Gruss, L. F., & Keil, A. (2015). Aversive learning shapes neuronal orientation tuning in human visual cortex. Nat Commun, 6, 7823. doi:10.1038/ncomms8823

Miskovic, V., & Keil, A. (2013). Visuocortical changes during delay and trace aversive conditioning: evidence from steady-state visual evoked potentials. Emotion, 13(3), 554–561. doi:10.1037/a0031323

Ohman, A., Flykt, A., & Esteves, F. (2001). Emotion drives attention: detecting the snake in the grass. Journal of experimental psychology. General, 130(3), 466–478.

Ohman, A., Lundqvist, D., & Esteves, F. (2001). The face in the crowd revisited: a threat advantage with schematic stimuli. J Pers Soc Psychol, 80(3), 381–396.

Peckham, A. D., McHugh, R. K., & Otto, M. W. (2010). A meta-analysis of the magnitude of biased attention in depression. Depression and anxiety, 27(12), 1135–1142. doi:10.1002/da.20755

Picardo, R., Baron, A. S., Anderson, A. K., & Todd, R. M. (2016). Tuning to the Positive: Age-Related Differences in Subjective Perception of Facial Emotion. PLoS ONE, 11(1), e0145643. doi:10.1371/journal.pone.0145643

Pourtois, G., Schettino, A., & Vuilleumier, P. (2013). Brain mechanisms for emotional influences on perception and attention: what is magic and what is not. Biological psychology, 92(3), 492–512. doi:10.1016/j.biopsycho.2012.02.007

Purcell, D. G., & Stewart, A. L. (2010). Still another confounded face in the crowd. Attention, perception & psychophysics, 72(8), 2115–2127. doi:10.3758/APP.72.8.2115

Rasch, B., Spalek, K., Buholzer, S., Luechinger, R., Boesiger, P., Papassotiropoulos, A., & de Quervain, D. J. (2009a). A genetic variation of the noradernergic system is related to differential amygdala activation during encoding of emotional memories. Proc Natl Acad Sci U S A.

Rasch, B., Spalek, K., Buholzer, S., Luechinger, R., Boesiger, P., Papassotiropoulos, A., & de Quervain, D. J. (2009b). A genetic variation of the noradrenergic system is related to differential amygdala activation during encoding of emotional memories. Proceedings of the National Academy of Sciences of the United States of America, 106(45), 19191–19196. doi:10.1073/pnas.0907425106

Rodriguez, S., Gaunt, T. R., & Day, I. N. (2009). Hardy-Weinberg equilibrium testing of biological ascertainment for Mendelian randomization studies. American journal of epidemiology, 169(4), 505–514. doi:10.1093/aje/kwn359

Roozendaal, B., Luyten, L., de Voogd, L. D., & Hermans, E. J. (2016). Importance of amygdala noradrenergic activity and large-scale neural networks in regulating emotional arousal effects on perception and memory. Behav Brain Sci, 39, e222. doi:10.1017/S0140525X15001934

Rossi, V., Vanlessen, N., Bayer, M., Grass, A., Pourtois, G., & Schacht, A. (2017). Motivational Salience Modulates Early Visual Cortex Responses across Task Sets. J Cogn Neurosci, 1–12. doi:10.1162/jocn_a_01093

Sara, S. J. (2009). The locus coeruleus and noradrenergic modulation of cognition. Nature Reviews Neuroscience, 10(3), 211–223. doi:10.1038/nrn2573

Sara, S. J., & Bouret, S. (2012). Orienting and reorienting: the locus coeruleus mediates cognition through arousal. Neuron, 76(1), 130–141. doi:10.1016/j.neuron.2012.09.011

Savage, R. A., Becker, S. I., & Lipp, O. V. (2016). Visual search for emotional expressions: Effect of stimulus set on anger and happiness superiority. Cognition & emotion, 30(4), 713–730. doi:10.1080/02699931.2015.1027663

Savage, R. A., Lipp, O. V., Craig, B. M., Becker, S. I., & Horstmann, G. (2013). In search of the emotional face: anger versus happiness superiority in visual search. Emotion, 13(4), 758–768. doi:10.1037/a0031970

Small, K. M., Brown, K. M., Forbes, S. L., & Liggett, S. B. (2001). Polymorphic deletion of three intracellular acidic residues of the alpha 2B-adrenergic receptor decreases G protein-coupled receptor kinase-mediated phosphorylation and desensitization. J Biol Chem, 276(7), 4917–4922. doi:10.1074/jbcM008118200[pii]

Stein, T., & Sterzer, P. (2012). Not just another face in the crowd: detecting emotional schematic faces during continuous flash suppression. Emotion, 12(5), 988–996. doi:10.1037/a0026944

Surguladze, S. A., Young, A. W., Senior, C., Brebion, G., Travis, M. J., & Phillips, M. L. (2004). Recognition accuracy and response bias to happy and sad facial expressions in patients with major depression. Neuropsychology, 18(2), 212–218. doi:10.1037/0894-4105.18.2.212

Todd, R. M., Cunningham, W. A., Anderson, A. K., & Thompson, E. (2012). Affect-biased attention as emotion regulation. Trends in Cognitive Sciences, 16(7), 365–372. doi:10.1016/j.tics.2012.06.003

Todd, R. M., Ehlers, M. R., Muller, D. J., Robertson, A., Palombo, D. J., Freeman, N., … Anderson, A. K. (2015). Neurogenetic variations in norepinephrine availability enhance perceptual vividness. J Neurosci, 35(16), 6506–6516. doi:10.1523/JNEUROSCI.4489-14.2015

Todd, R. M., Evans, J. W., Morris, D., Lewis, M. D., & Taylor, M. J. (2011). The changing face of emotion: age-related patterns of amygdala activation to salient faces. Soc Cogn Affect Neurosci, 6(1), 12–23. doi:nsq007[pii]10.1093/scan/nsq007

Todd, R. M., & Manaligod, M. G. M. (2017). Implicit guidance of attention: The priority state space framework. Cortex; a journal devoted to the study of the nervous system and behavior. doi:10.1016/j.cortex.2017.08.001

Todd, R. M., Muller, D. J., Lee, D. H., Robertson, A., Eaton, T., Freeman, N., … Anderson, A. K. (2013). Genes for emotion-enhanced remembering are linked to enhanced perceiving. Psychol Sci, 24(11), 2244–2253. doi:10.1177/0956797613492423[pii]

Todd, R. M., Muller, D. J., Palombo, D. J., Robertson, A., Eaton, T., Freeman, N., … Anderson, A. K. (2014). Deletion variant in the ADRA2B gene increases coupling between emotional responses at encoding and later retrieval of emotional memories. Neurobiol Learn Mem, 112, 222–229. doi:10.1016/j.nlm.2013.10.008

Todd, R. M., Talmi, D., Schmitz, T. W., Susskind, J., & Anderson, A. K. (2012). Psychophysical and neural evidence for emotion-enhanced perceptual vividness. J Neurosci, 32(33), 11201–11212. doi:10.1523/JNEUROSCI.0155-12.2012

Walther, D., & Koch, C. (2006). Modeling attention to salient proto-objects. Neural Netw, 19(9), 1395–1407. doi:S0893-6080(06)00215-2[pii]10.1016/j.neunet.2006.10.001

Waterhouse, B. D., & Navarra, R. L. (2018). The locus coeruleus-norepinephrine system and sensory signal processing: A historical review and current perspectives. Brain Res. doi:10.1016/j.brainres.2018.08.032

West, G. L., Anderson, A. A., & Pratt, J. (2009). Motivationally significant stimuli show visual prior entry: evidence for attentional capture. Journal of experimental psychology. Human perception and performance, 35(4), 1032–1042. doi:10.1037/a0014493

Wieser, M. J., & Keil, A. (2011). Temporal Trade-Off Effects in Sustained Attention: Dynamics in Visual Cortex Predict the Target Detection Performance during Distraction. The Journal of neuroscience: the official journal of the Society for Neuroscience, 31(21), 7784–7790. doi:10.1523/JNEUROSCI.5632-10.2011

Wieser, M. J., McTeague, L. M., & Keil, A. (2012). Competition effects of threatening faces in social anxiety. Emotion, 12(5), 1050–1060. doi:10.1037/a0027069

Wieser, M. J., Reicherts, P., Juravle, G., & von Leupoldt, A. (2016). Attention mechanisms during predictable and unpredictable threat - A steady-state visual evoked potential approach. Neuroimage, 139, 167–175. doi:10.1016/j.neuroimage.2016.06.026

Xie, W., Cappiello, M., Meng, M., Rosenthal, R., & Zhang, W. (2018). ADRA2B deletion variant and enhanced cognitive processing of emotional information: A meta-analytical review. Neurosci Biobehav Rev, 92, 402–416. doi:10.1016/j.neubiorev.2018.05.010

