## Supplemental correlation plots for "Affectively biased competition: sustained attention is tuned to rewarding expressions and is not modulated by norepinephrine receptor gene variant"

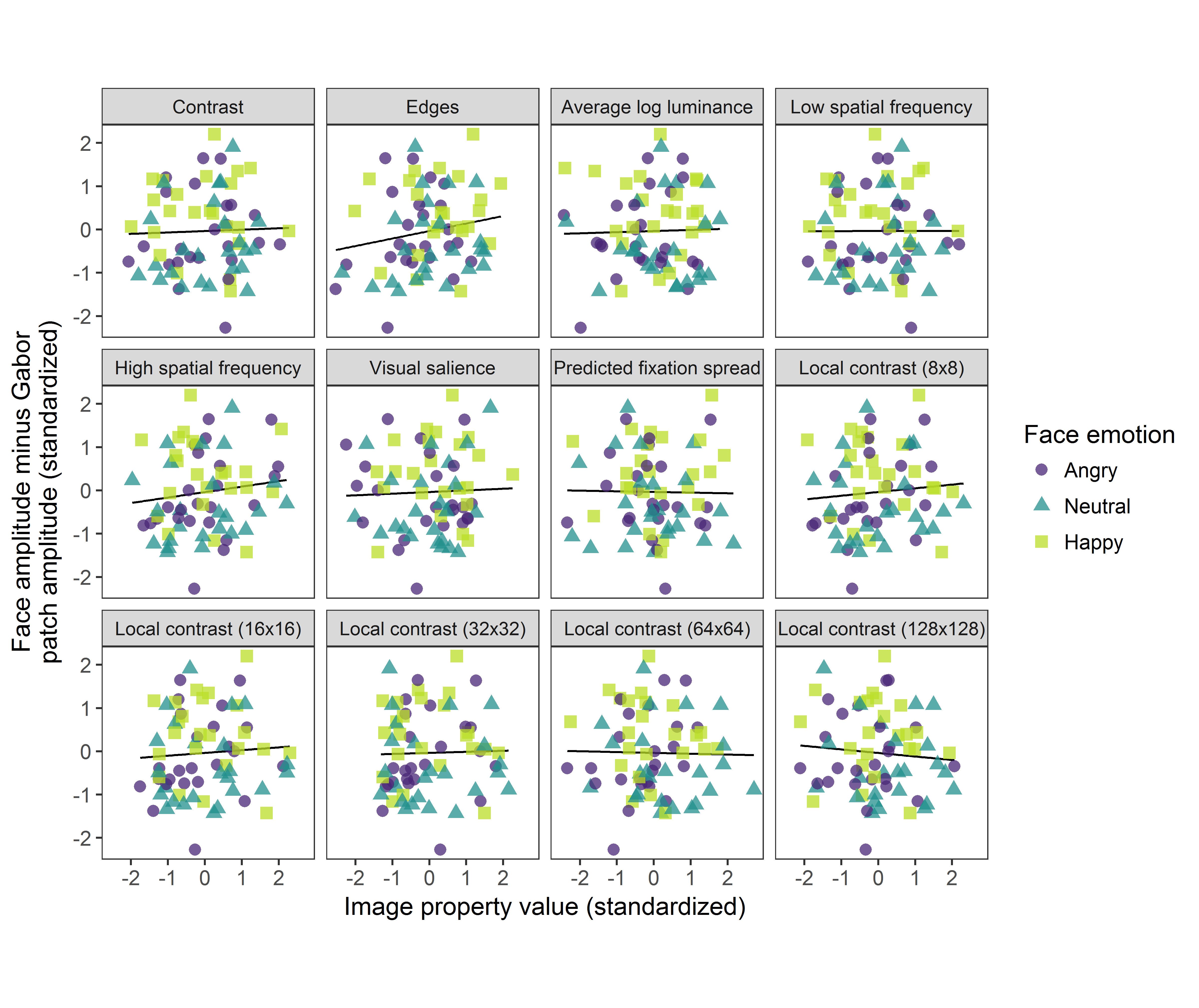

**Supplementary Figure 1**. Correlations between metrics of low level visual features and ssVEP difference scores for trials with happy, angry and neutral faces. Edges modestly correlated with ssVEP difference scores, but this correlation was not significant. All other low level visual features did not correlate with ssVEP differences. The happy face competition advantage cannot be explained by differences in these measured low level visual features.
